# Magnetoactive hydrogels to probe curvature-directed endothelial cell mechanosensing

**DOI:** 10.64898/2026.05.04.722723

**Authors:** Avinava Roy, Grace K Hinds, Joshua Yu-Chung Liu, Rishab Yanala, Arina Velieva, Claudia Loebel

## Abstract

The vascular system exhibits complex, non-planar geometries that become further distorted during pathological remodeling, including arterial tortuosity and aneurysms. Although hemodynamic shear stress is a well-established regulator of vascular function, the direct effects of curvature as an intrinsic geometric cue remain poorly defined. This is largely because existing *in vitro* models are static and fail to capture the dynamic changes that accompany disease progression. To address this gap, we used a magnetoactive hydrogel platform that enables real-time, on-demand curvature of endothelial monolayers to reproduce clinically established tortuosity metrics. Using this system, we found that elevated curvature increased nuclear localization of yes-associated protein (YAP), with the strongest response in convex relative to concave regions of highly tortuous endothelial monolayers. This mechanosensitive response was accompanied by reduced VE-Cadherin junctional thickness and increased membrane localization of endothelial nitric oxide synthase. Together, these findings identify local curvature, independent of shear stress, as a regulator of endothelial cell mechanosensing and function, and establish a dynamic hydrogel platform for isolating geometric regulation from shear stress inputs in vascular mechanobiology.

## 1. Introduction

Tubular biological structures inside the human body show complex and heterogeneous geometries that shape the mechanical environment experienced by resident cells.[1] In the vasculature, these geometries range from the high curvature of microcapillaries to the branching architecture of large arteries and become further distorted in disease states such as atherosclerosis[2,3] and aneurysm formation[4,5]. Such pathological remodeling often gives rise to tortuous or abnormally curved vessel segments that are associated with endothelial dysfunction, including increased permeability that may lead to inflammatory activation.[6] Although blood flow-derived shear stress is widely recognized as a central regulator of endothelial cell function, it is itself influenced by vessel geometry.[7] This relationship makes it difficult to separate the effects of flow from those of curvature alone.

Most studies of endothelial mechanobiology have therefore focused on shear stress to explain local vascular dysfunction.[8] However, in vessels with changes in geometry or tortuous morphology, endothelial cells are also subjected to curvature-dependent mechanical deformation that may act independently of flow.[9] This raises the possibility that local curvature serves as a primary mechanosensitive cue that primes endothelial cell dysfunction before, or in parallel with, hemodynamic changes.[10] Despite this potential importance, the role of changes in curvature itself has remained relatively unexplored, in part because of the difficulty to introduce spatiotemporal changes in localized curvature onto endothelial cells while decoupling from shear stress.

Several *in vitro* studies have incorporated aspects of vascular geometry using curved substrates, microchannels,[11] engineered stenoses, aneurysms, bifurcations,[12] or perfusable vessel models. These studies have provided evidence that geometry directs changes in endothelial cell remodeling, migration and cytoskeletal structure.[10] However, in most cases, the observed effects were coupled to the flow conditions used to culture or stimulate the cells. This limits interpretation of curvature as an independent cue. Moreover, few platforms have enabled the analysis of planar-to-curved transitions within a continuous endothelial monolayer, where localized phenotypic differences may emerge. Even fewer studies have investigated dynamic development of curvature, despite the fact that vascular remodeling occurs over time and likely engages mechanosensing pathways.[13–15]

Here, we build upon our previously developed magnetoactive hydrogel platform[16] to dynamically introduce curvature to endothelial monolayers and investigate endothelial cell mechanosensing and function. Using this system, we show high tortuosity acts a mechanical cue that promotes nuclear translocation of Yes-associated protein (YAP), reduces thickness of vascular endothelial cadherin (VE-Cadherin) junctions, and alters endothelial nitric oxide synthase (eNOS) localization. These effects were particularly pronounced in regions of convex curvature. Together, these findings identify local substrate curvature, independent of shear stress, as a driver of endothelial cell functional changes and establish dynamic curvature control as a useful approach for studying vascular mechanobiology.

## 2. Results and discussion

### 2.1 Flat actuation of programmed magnetic hydrogels induces limited endothelial cell responses

To fabricate magnetic hydrogels, we incorporated 150 wt.% ferromagnetic neodymium iron (NdFe) microparticles into alginate/polyacrylamide double network hydrogels.[16] In the unprogrammed state, the embedded NdFe particles possess randomly oriented magnetic dipoles and therefore net zero magnetic moment (Fig. 1A(i)). After pressing the hydrogels between flat molds and exposing them to a 1 Tesla external magnetic field, the particles acquired a net magnetic moment that was retained after field removal, resulting in hydrogels with a programmed magnetic field (Fig. 1A (ii-iii)). This programmed state enabled subsequent actuation with substantially lower magnetic fields (Fig. 1A(iv)).

**Fig. 1.**
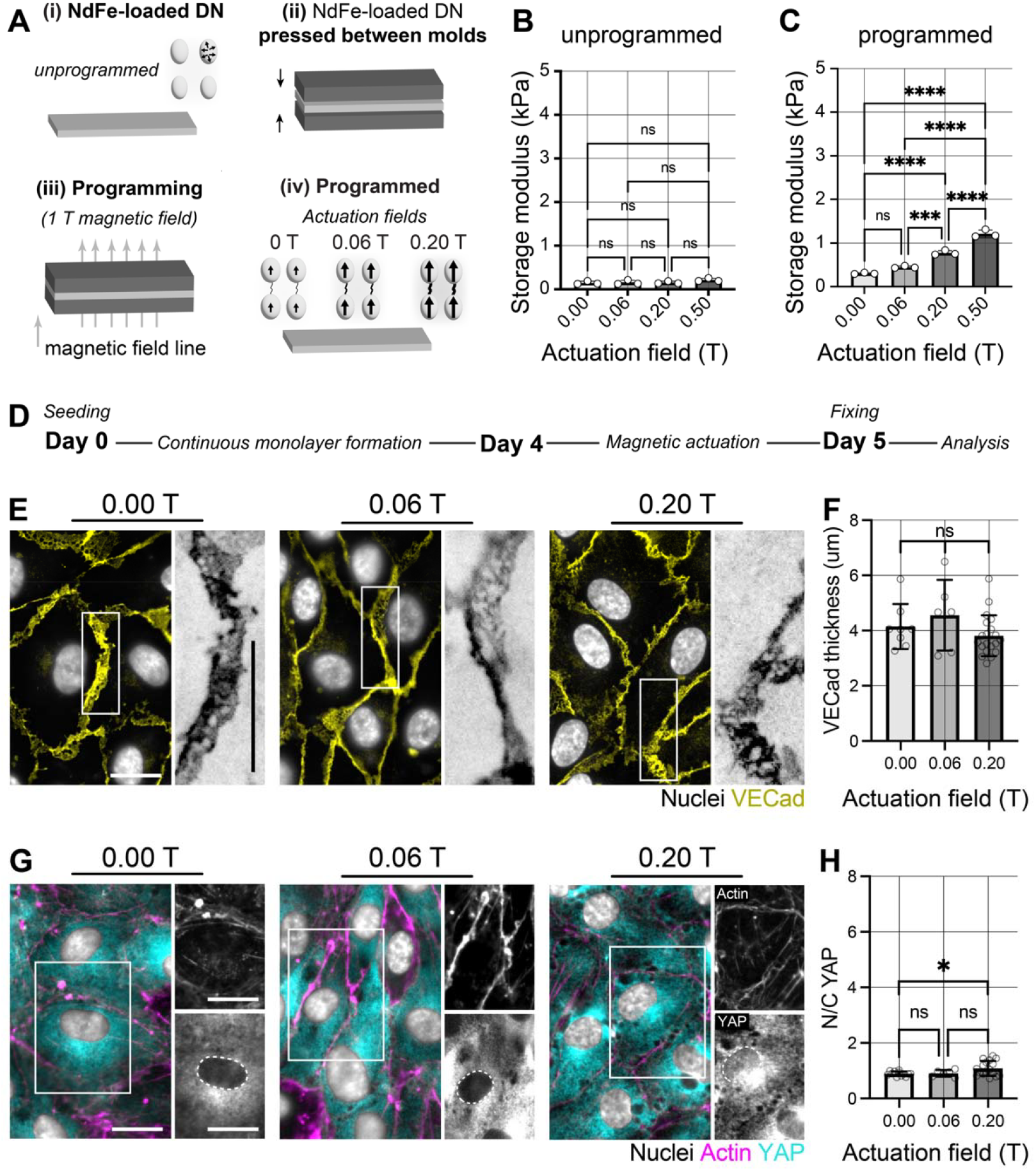
Flat magnetic hydrogels induce limited endothelial cell response. **A** Schematics illustrating the steps of programming (i) unprogrammed NdFe-loaded double network hydrogels containing particles with random magnetic dipoles by (ii) sandwiching hydrogels between two flat resin molds, followed by (iii) exposure to a high-intensity 1 T programming field, which (iv) enables a non-random magnetic dipole moment (increasing with actuation field (0.00 T to 0.20 T) of NdFe particles embedded inside the hydrogel. **B** Shear storage moduli of unprogrammed magnetic hydrogels as a function of actuation field strength (n = 3 hydrogels per group from 3 independent experiments). **C** Shear storage moduli of programmed magnetic hydrogels as a function of actuation field strength (n = 3 hydrogels per group). **D** Experimental timeline showing steps of seeding, monolayer formation, magnetic actuation, fixing and finally, analysis of human umbilical vein endothelial cells (hUVECs). **E** Representative fluorescent images of nuclei (Hoechst) and VE-Cadherin of hUVEC monolayer cultured atop programmed magnetic hydrogels exposed to actuation fields of 0.00 T, 0.06 T and 0.20 T for 24 h (scale bars 20 μm). **F** Quantification of VE-Cadherin junction thickness between adjacent cells (0.00 T: n = 8 ROIs; each ROI = 100 μm^2^; N = 2, 0.06 T: n = 6 ROIs; each ROI = 100 μm^2^; N = 2, 0.20 T: n = 20 ROIs; each ROI = 100 μm^2^; N = 2). **G** Representative fluorescent images of nuclei (Hoechst), F-actin, and Yes-Associated Protein (YAP) of hUVEC monolayer atop programmed magnetic hydrogels exposed to actuation fields of 0.00 T, 0.06 T, and 0.20 T for 24 h (scale bars 20 μm). **H** Quantification of YAP nuclear-to-cytoplasmic (N/C) ratio of mean fluorescence intensities (0.00 T: n = 12 ROIs; each ROI = 100 μm^2^; N = 3, 0.06 T: n = 6 ROIs; each ROI = 100 μm^2^; N = 2, 0.20 T: n = 17 ROIs; each ROI = 100 μm^2^; N = 3). **B-H** N = independent experiments, ****p < 0.0001, ***p < 0.001, *p < 0.05, ns: not significant by one-way ANOVA with Bonferroni post hoc testing.

Shear rheology of unprogrammed magnetic hydrogels demonstrated little change in shear modulus with increasing actuation fields up to 0.50 T (Fig. 1B). In contrast, magnetic hydrogels post-programming showed a field-dependent increase in shear modulus, rising from 0.30 ± 0.02 kPa at 0.00 T to 1.21 ± 0.09 kPa at 0.50 T (Fig. 1A). Programming itself induced a two-fold increase in the baseline modulus (Fig. S1, Supporting Information). As actuation at or above 0.5 T may re-program the dipoles, we chose to continue with 0.06 and 0.2 T actuation for subsequent studies.[17]

We next asked whether endothelial cells respond to these actuation-induced changes in substrate modulus. Endothelial cells were seeded on programmed hydrogels and cultured for 4 days to form a continuous monolayer before exposure to actuation fields of 0.06 and 0.2 T for 24 h (Fig. 1D). Staining for VE-Cadherin revealed junctions between all cells and across all conditions (Fig. 1E). Quantification of average thickness did not identify significant differences between groups (Fig. 1F). This suggests that actuation-induced changes in hydrogel modulus had little effect on gross junctional morphology over this time frame. In contrast, cells cultured on 0.00 and 0.06 T hydrogels primarily showed cortical actin, whereas cells on 0.20 T hydrogels had more prominent actin fibers spanning the cell body. Consistent with this shift, immunofluorescence staining for YAP, an established marker of mechanical activation for endothelial cells,[18,19] showed mostly cytoplasmic localization under 0.00 and 0.06 T conditions, while cells on 0.20 T hydrogels showed some nuclear enrichment (Fig. 1G). Quantification of the nuclear-to-cytoplasmic (N/C) YAP ratio showed a modest increase in YAP nuclear translocation at 0.02 T actuation field (Fig. 1H). Together, these data indicate that magnetic actuation of flat programmed hydrogels induces only limited endothelial responses, with modest actin reorganization and slight YAP nuclear enrichment but little change in VE-Cadherin junction thickness. These findings establish the flat hydrogel condition as an important baseline and suggest that changes in substrate modulus alone are insufficient to drive endothelial remodeling.[20] It further highlights the need to investigate additional substrate-mediated cues that may regulate endothelial phenotype and function.

### 2.2 Programming magnetic hydrogels to mimic arterial tortuosity

Having established that magnetic actuation of flat programmed hydrogels elicits only limited endothelial cell responses, we next asked how dynamic changes in hydrogel curvature modulate endothelial function. To this end, we used our previously established resin mold-based protocol for programmable patterning.[16] Unprogrammed magnetic hydrogels were pressed between two patterned resin molds and exposed to a 1 T magnetic field to spatially program the magnetic moments of the embedded particles according to the mold architecture (Fig. 2A(i-ii)). After removal from the molds, the physical crosslinks of the magnetic hydrogels enabled flattening in the absence of a field and fully reversible recovery of the programmed pattern upon exposure to a low-intensity actuation field of 0.2 T (Fig. 2A(iii)).

**Fig. 2.**
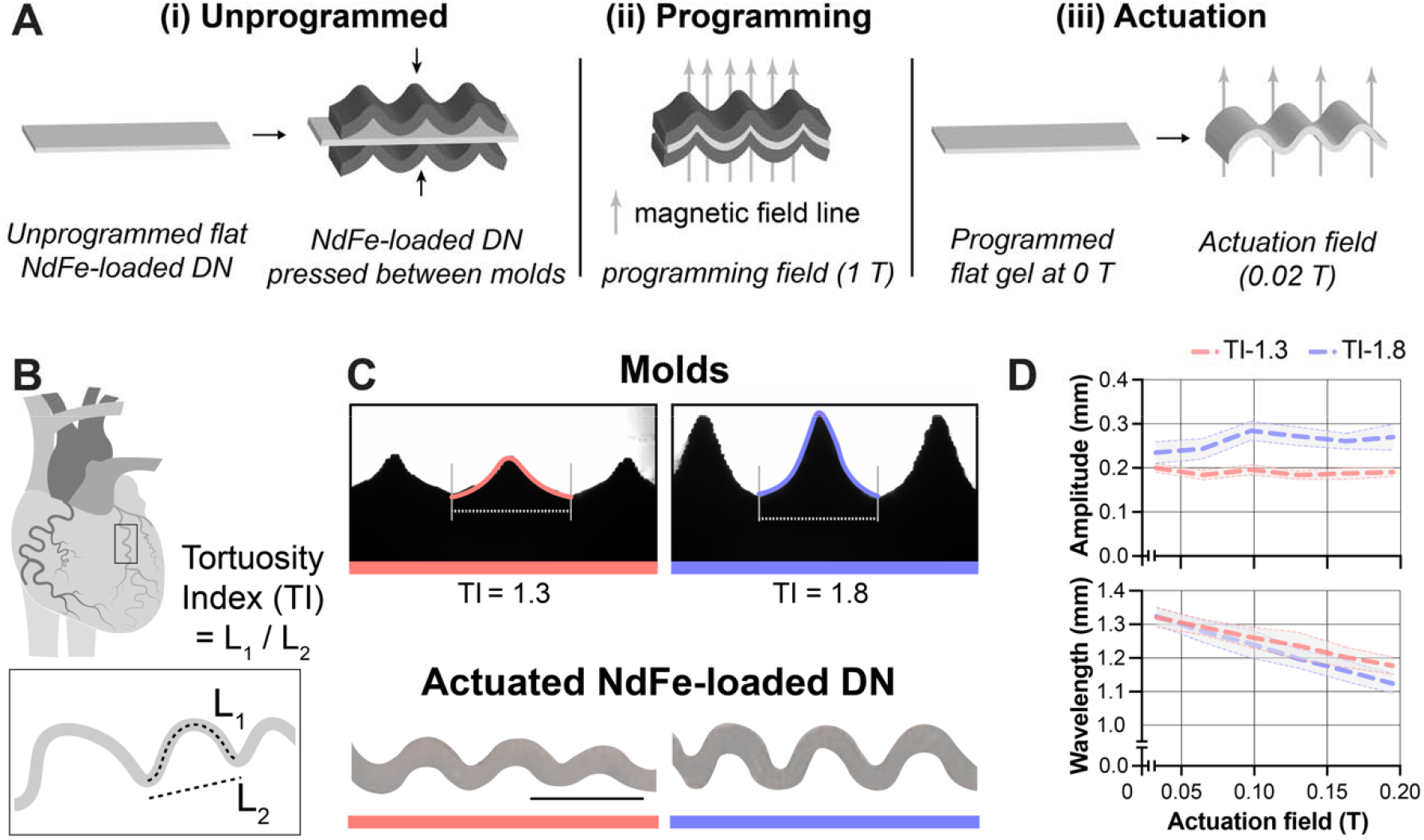
Programming magnetic hydrogels to mimic arterial tortuosity. **A** Schematics illustrating the patterning of magnetic hydrogels by (i) sandwiching hydrogels between two resin molds, followed by (ii) exposure to a high-intensity 1 T programming field, which (iii) enables reversible patterning using low-intensity 0.02 T actuation field. **B** Schematic illustrating coronary arteries with varying tortuosity index (TI) calculated as the ratio of curved (L_1_) to straight (L_2_) end-to-end lengths of one artery. **C** Representative images of resin molds (top) to obtain hydrogel patterns (bottom) that mimic the tortuosity of TI 1.3 and TI 1.8, scale bar 1 mm). **D** Quantification of height and width of TI 1.3 and TI 1.8 hydrogel patterns as a function of the actuation field (n = 4 hydrogels from 4 independent experiments; shaded areas indicate mean ± s.d.).

As resin molds are tunable in their dimensions, we next designed molds to mimic curvatures with selected tortuosity indices (TI), defined as the ratio of the curved length (L_1_) to the straight end-to-end length (L_2_, Fig. 2B). Clinically relevant TI values of 1.3 and 1.8 represent low and high tortuosity indices.[21,22] Accordingly, we fabricated molds with L_1_/L_2_ values of 1.29/1.05 mm (TI 1.3) and 1.89/1.05 mm (TI 1.8), which generated patterns resembling tortuous human arteries (Fig. 2C). Quantification of the hydrogel shape as a function of actuation field showed an increase in pattern height and a decrease in width for both TI 1.3 and TI 1.8 hydrogels (Fig. 2D). Notably, the height of TI 1.3 hydrogels plateaued at 0.05 T, whereas TI 1.8 hydrogels continued to increase in height up to 0.10 T. In contrast, pattern width continued to decrease up to 0.20 T, likely reflecting increasing compression of the programmed structure as the magnetic strength increased. Together, these results show that programming magnetic hydrogels within 3D-printed resin molds enables on-demand formation of patterns that mimic clinically relevant vascular tortuosity.

### 2.3 Local hydrogel curvature promotes mechanical activation of endothelial monolayers

Since we observed magnetic actuation of hydrogels in the flat state induces little endothelial remodeling, we next investigated whether local curvature more strongly regulates endothelial mechanical activation. Endothelial cells allowed to form a monolayer on initially flat hydrogels were then exposed to a 0.20 T actuation field to evolve into flat (TI 1.0), moderately tortuous (TI 1.3) and highly tortuous (TI 1.8) patterns. Within 12 h of curvature induction, cells on TI 1.3 and TI 1.8 hydrogels showed less cortical actin and more fibers spanning the cell body, compared to flat TI 1.0 hydrogels (Fig. 3A). Consistent with this shift, immunofluorescence staining for YAP showed increased nuclear localization in cells cultured on both TI 1.3 and TI 1.8 hydrogel patterns (Fig. 3A). Quantification of the N/C YAP ratio confirmed an increase from 1.22 ± 0.33 (TI 1.0) to 1.45 ± 0.31 (TI 1.3) and 1.85 ± 0.60 (TI 1.8) (Fig. 3B).

**Fig. 3.**
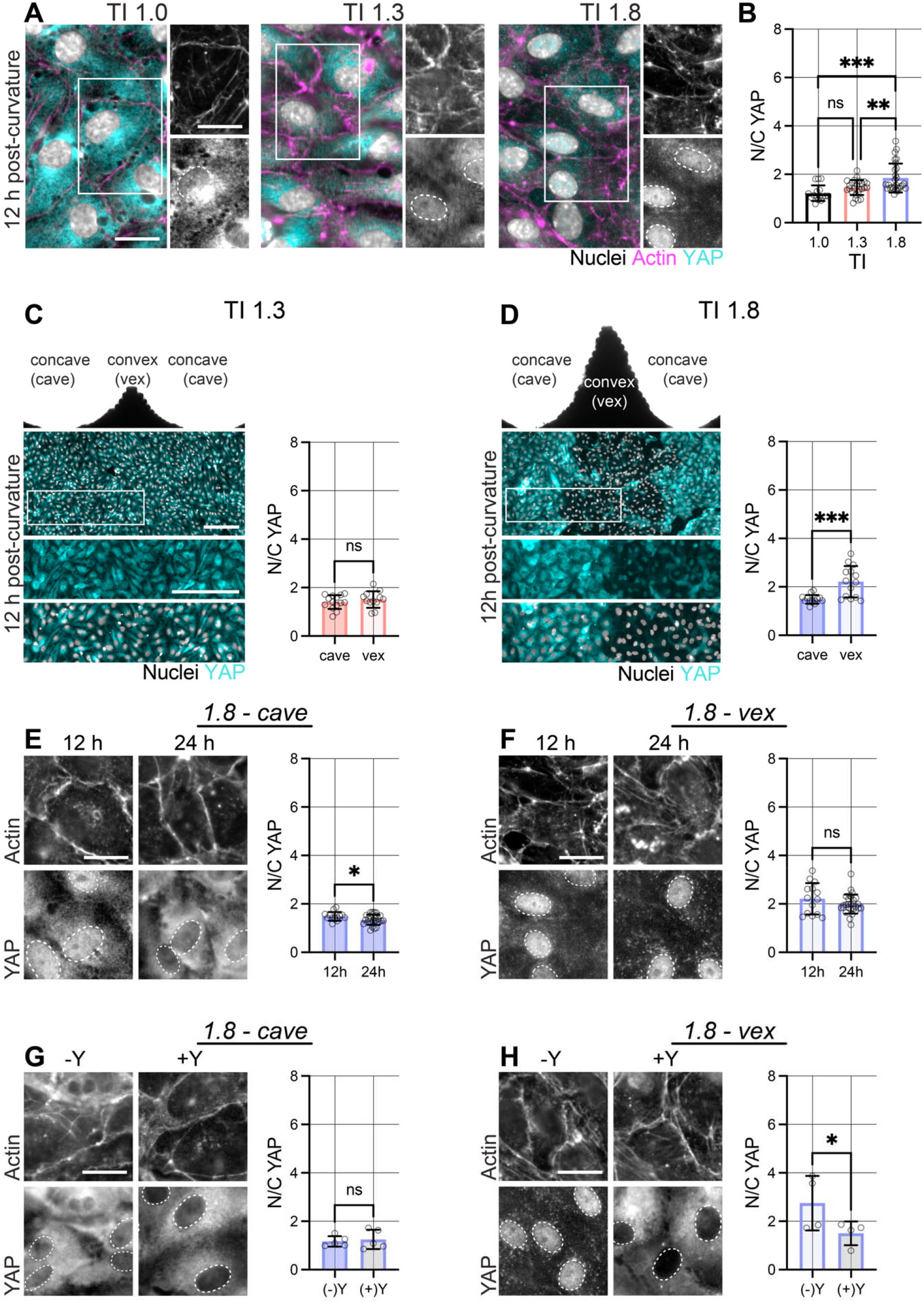
Local convex curvature increases endothelial YAP nuclear localization. **A** Representative fluorescent images of nuclei (Hoechst), actin cytoskeleton and YAP of endothelial cells cultured atop programmed TI 1.0 (flat), TI 1.3, and TI 1.8 hydrogels at 12 h after induction of curvature. (0.2 T actuation field, scale bars 20 μm). **B** Quantification of YAP nuclear-to-cytoplasmic (N/C) ratio of mean fluorescence intensities across the entire sample surface (flat: n = 16 ROIs; each ROI = 100 μm^2^; N = 3, TI 1.3: n = 26 ROIs; each ROI = 100 μm^2^; N = 6, TI 1.8: n = 28 ROIs; each ROI = 100 μm^2^; N = 6). **C** Representative mold image atop with fluorescent images and quantification of local YAP N/C ratio of endothelial cells on TI 1.3 convex (vex) and concave (cave) regions at 12 h (scale bars 20 μm, cave: n = 14 ROIs; each ROI = 100 μm^2^; N = 3, vex: n = 12 ROIs; each ROI = 100 μm^2^; N = 3). **D** Representative mold image atop with fluorescent images and quantification of local YAP N/C ratio of endothelial cells atop TI 1.8 convex (vex) and concave (cave) regions at 12 h (scale bars 20 μm, cave: n = 14 ROIs; each ROI = 100 μm^2^; N = 3, vex: n = 14 ROIs; each ROI = 100 μm^2^; N = 3). **E** Representative fluorescent images of F-actin and YAP (bottom) and quantification of YAP N/C ratios at 12 h and 24 h atop concave (cave) regions (scale bars 20 μm, 12h: n = 14 ROIs; each ROI = 100 μm^2^; N = 3, 24h: n = 27 ROIs; each ROI = 100 μm^2^; N = 5). **F** Representative fluorescent images of F-actin and YAP (bottom) and quantification of YAP N/C ratios at 12 h and 24 h atop convex (vex) regions (scale bars 20 μm, 12h: n = 14 ROIs; each ROI = 100 μm^2^; N = 3, 24h: n = 28 ROIs; each ROI = 100 μm^2^; N = 5) **G** Representative images of F-actin and YAP with quantification of YAP N/C ratio without and with Y-27 (Y, 10 μM) for 24 h atop concave (cave) regions (scale bars 20 μm, (−)Y: n = 5 ROIs; each ROI = 100 μm^2^; N = 2, (+)Y: n = 5 ROIs; each ROI = 100 μm^2^; N = 2). **H** Representative images of F-actin and YAP with quantification of YAP N/C ratios without and with Y-27 (Y, 10 μM) for 24 h atop convex (vex) regions (scale bars 20 μm, (−)Y: n = 7 ROIs; each ROI = 100 μm^2^; N = 2, (+)Y: n = 4 ROIs; each ROI = 100 μm^2^; N = 2). **B-H** N = independent experiments, ***p < 0.001, **p < 0.01, *p < 0.05, ns: not significant by two-tailed Student’s *t*-tests (two experimental groups) and one-way analysis of variance (ANOVA) with Bonferroni post hoc testing (more than two experimental groups).

The high variability of N/C YAP ratios observed on TI 1.8 hydrogels prompted us to investigate whether local curvature differentially regulates endothelial mechanosensing. Tile-scans spanning multiple ROIs of cells on TI 1.3 hydrogels showed relatively similar YAP localization across concave and convex regions, which was supported by similar N/C YAP ratios in concave and convex regions (Fig. 3C). In contrast, cells on TI 1.8 hydrogels displayed region-dependent YAP localization with a 1.5-fold higher N/C YAP ratio in convex regions than in concave regions (Fig. 3D). These findings indicate that local convex curvature within highly tortuous hydrogels promotes endothelial mechanical activation.

We next asked whether curvature-induced changes in F-actin organization and YAP localization were sustained over time by extending culture for an additional 12 h. In concave regions, F-actin organization appeared similar at 12 and 24 h, although N/C YAP ratios were slightly reduced at 24 h (Fig. 3E). In contrast, cells in convex regions showed persisting prominent F-actin fibers spanning the cell body and higher YAP nuclear localization at both the 12 h and 24 h timepoints (Fig. 3F). To further probe whether intracellular contractility contributes to curvature-induced YAP activation, we treated cells with the Rho-associated protein kinase inhibitor Y27632 (Y27) at the onset of curvature induction and quantified local N/C YAP ratios after 24 h. In concave regions, cortical F-actin and predominantly cytoplasmic YAP were observed and remained largely unchanged with Y27 treatment (Fig. 3G). In contrast, we mostly observed cytoplasmic YAP localization in Y27 treated cells on convex regions, thereby having reduced N/C YAP ratios (Fig. 3H). Moreover, the N/C YAP levels of the cells atop convex regions of the Y27 treated samples were comparable to those measured in concave regions (Fig. S2, Supporting Information). These results suggest that convex curvature promotes F-actin reorganization and YAP nuclear translocation in a Rho-associated protein kinase-dependent manner.

### 2.4 Local convex curvature induces sustained thinning of VE-Cadherin junctions

Given the local changes in YAP nuclear localization on TI 1.8 hydrogels, we next asked whether local curvature also alters cell-cell junctions, a vital component of the endothelial monolayer. To visualize cell-to-cell junction remodeling, we stained for VE-Cadherin 12 h after curvature induction. Inverted grayscale images showed thinner VE-Cadherin junctions between cells on both concave and convex regions compared to flat controls (Fig. 4A). Quantification confirmed a reduction in junction thickness from 3.50 ± 0.35 μm on flat samples to 2.85 ± 0.52 μm in concave regions and 2.66 ± 0.38 μm in convex regions (Fig. 4B). This observation suggests that curvature-induced endothelial activation is accompanied by junctional thinning. To determine whether these changes were sustained over time, we quantified VE-Cadherin thickness after an additional 12 h of culture. Interestingly, junction thickness of cells atop concave regions increased after the additional 12 h such that there was little difference in junctional thickness between cells experiencing flat vs concave regions by 24 h post onset of curvature (Fig. S3, Supporting Information). Junctional thickness between cells on convex regions however, remained largely unchanged between 12 and 24 h (Fig. 4C), which is consistent with the persistence of elevated YAP nuclear localization over this time frame (Fig. 3E, F).

**Fig. 4.**
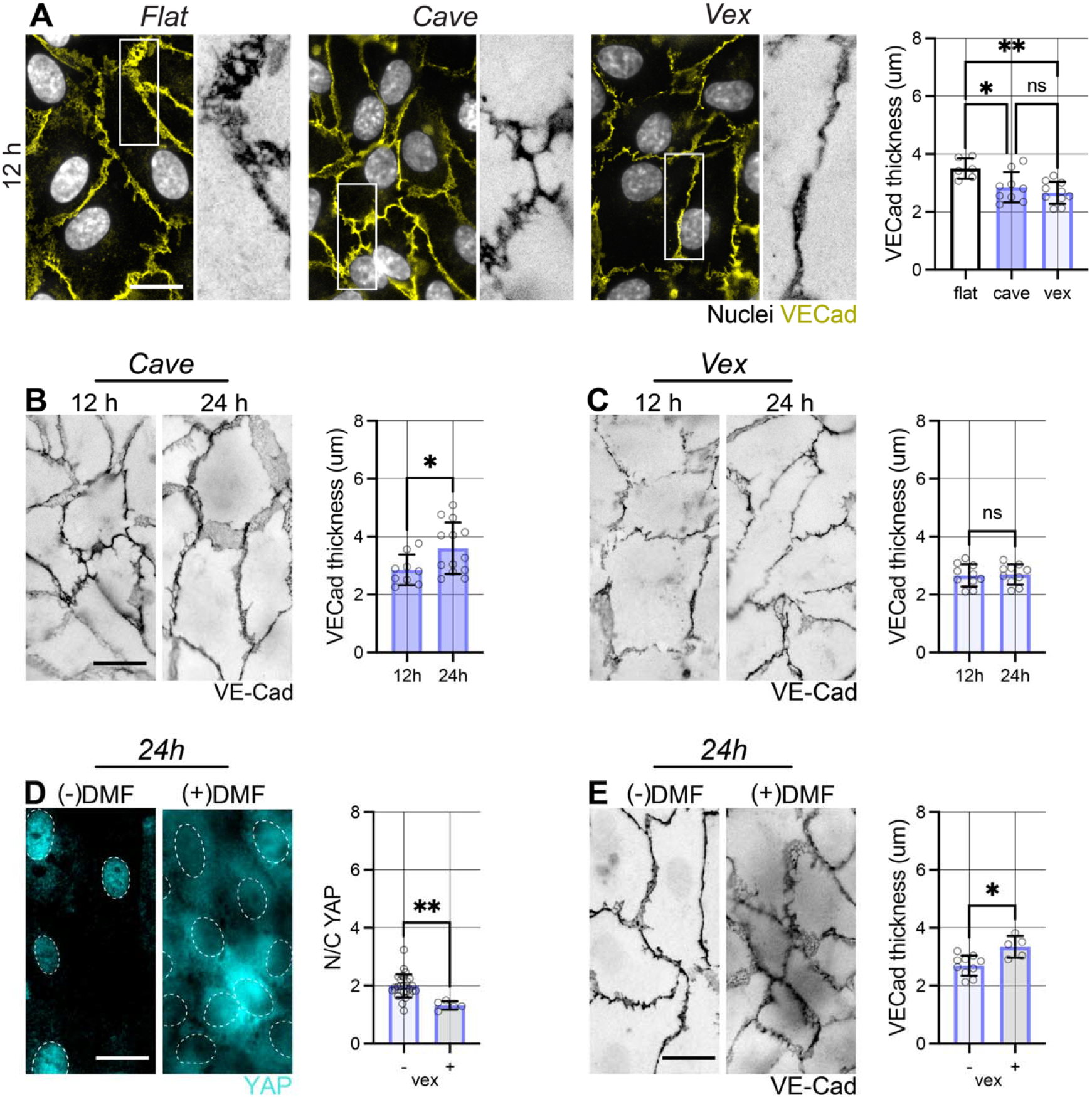
Convex curvature promotes VE-Cadherin junction thinning. **A** Representative fluorescent images of nuclei (Hoechst) and VE Cadherin, with inverted grayscale views of insets and quantification of VE-Cadherin thickness of endothelial cells atop programmed flat and TI 1.8 concave (cave) and convex (vex) regions of magnetic hydrogels at 12 h (scale bars 20 μm, flat: n = 6 ROIs; each ROI = 100 μm^2^; N = 2, cave: n = 9 ROIs; each ROI = 100 μm^2^; N = 3, vex: n = 10 ROIs; each ROI = 100 μm^2^; N = 3). **B** Representative inverted grayscale images and quantification of VE-Cadherin thickness at 12 h and 24 h atop concave regions of curvature (scale bars 20 μm, 12h: n = 9 ROIs; each ROI = 100 μm^2^; N = 3, 24h: n = 13 ROIs; each ROI = 100 μm^2^; N = 4). **C** Representative inverted grayscale images and quantification of VE-Cadherin thickness at 12 h and 24 h atop convex regions of curvature (scale bars 20 μm, 12 h: n = 10 ROIs; each ROI = 100 μm^2^; N = 3, 24 h: n = 10 ROIs; each ROI = 100 μm^2^; N = 3). **D** Representative fluorescent images and quantification of YAP N/C ratios on TI 1.8 convex regions without ((−) DMF) and with dimethyl fumarate ((+)DMF) treatment (50 μM) at 24 h (scale bars 20 μm, (−)DMF: n = 13 ROIs; each ROI = 100 μm^2^; N = 5, (+)DMF: n = 5 ROIs; each ROI = 100 μm^2^; N = 2). **E** Representative inverted grayscale images and quantification of VE-Cad thickness on TI 1.8 convex regions without ((−) DMF) and with dimethyl fumarate ((+)DMF) treatment (50 μM) at 24 h (scale bars 20 μm, (−)DMF: n = 10 ROIs; each ROI = 100 μm^2^; N = 3, (+)DMF:; each ROI = 100 μm^2^; N = 2). **A-E** N = independent experiments, **p < 0.01, *p < 0.05, ns: not significant by two-tailed Student’s *t*-tests (two experimental groups) and one-way analysis of variance (ANOVA) with Bonferroni post hoc testing (more than two experimental groups).

Therefore, we next examined whether YAP nuclear translocation contributes to junction thinning in convex regions. Dimethyl fumarate (DMF), which has been reported to inhibit YAP nuclear translocation without directly inhibiting cell mechanosensing,[23] was added at the onset of curvature induction, and cells were stained for both YAP and VE-Cadherin after 24 h. As expected, DMF reduced nuclear YAP staining in cells atop convex regions, which was confirmed by a decrease in N/C YAP ratio (Fig. 4C). Notably, DMF also increased endothelial junction thickness from 2.69 ± 0.11 μm in untreated to 3.34 ± 0.17 μm in DMF-treated samples (Fig. 4D), reaching similar thickness as observed for cells on concave regions (Fig. S4, Supporting Information). These findings suggest that inhibiting YAP nuclear translocation may rescue VE-Cadherin junctions from curvature-induced thinning. This is consistent with prior *in vitro* studies showing that, in confluent endothelial monolayers, YAP associates with VE-Cadherin-based adherens junctions which promotes cytoplasmic sequestration of YAP.[24] Previous studies have further suggested reduced endothelial barrier function in response to mechanical stretch (extensive convex curvature *in vitro*[25–27] and in vessel bifurcations *in vivo*[28]. Together, these results indicate that curvature-induced YAP activation is functionally linked to VE-Cadherin junction remodeling in endothelial monolayers.

### 2.5 Local hydrogel curvature regulates intracellular localization of endothelial nitric oxide synthase

Given that local curvature remodeled VE-Cadherin junctions, we next asked whether it also affects endothelial functional markers, focusing on the subcellular localization of endothelial nitric oxide synthase (eNOS), a key regulator of nitric oxide production and vascular homeostasis. Since eNOS activity is closely linked to its intracellular localization,[29] we performed immunofluorescence staining to visualize eNOS distribution in endothelial monolayers.

In flat controls and in concave regions of TI 1.8 hydrogels, eNOS was primarily localized to perinuclear regions 24 h after curvature induction. In contrast, in convex regions, eNOS was enriched along cell boundaries marked by VE-Cadherin-positive junctions indicated with the red triangles in the grayscale view (Fig. 5A). Quantification of membrane-to-cytoplasmic eNOS ratio confirmed this shift in intracellular localization (Fig. 5B). Previous studies have shown that sustained mechanical stretch promotes eNOS activation[30] and that plasma membrane localization is associated with changes in nitric oxide release[31]. Thus, the membrane enrichment of eNOS in convex regions is consistent with the notion that local curvature induces a functionally distinct endothelial state.

**Fig. 5.**
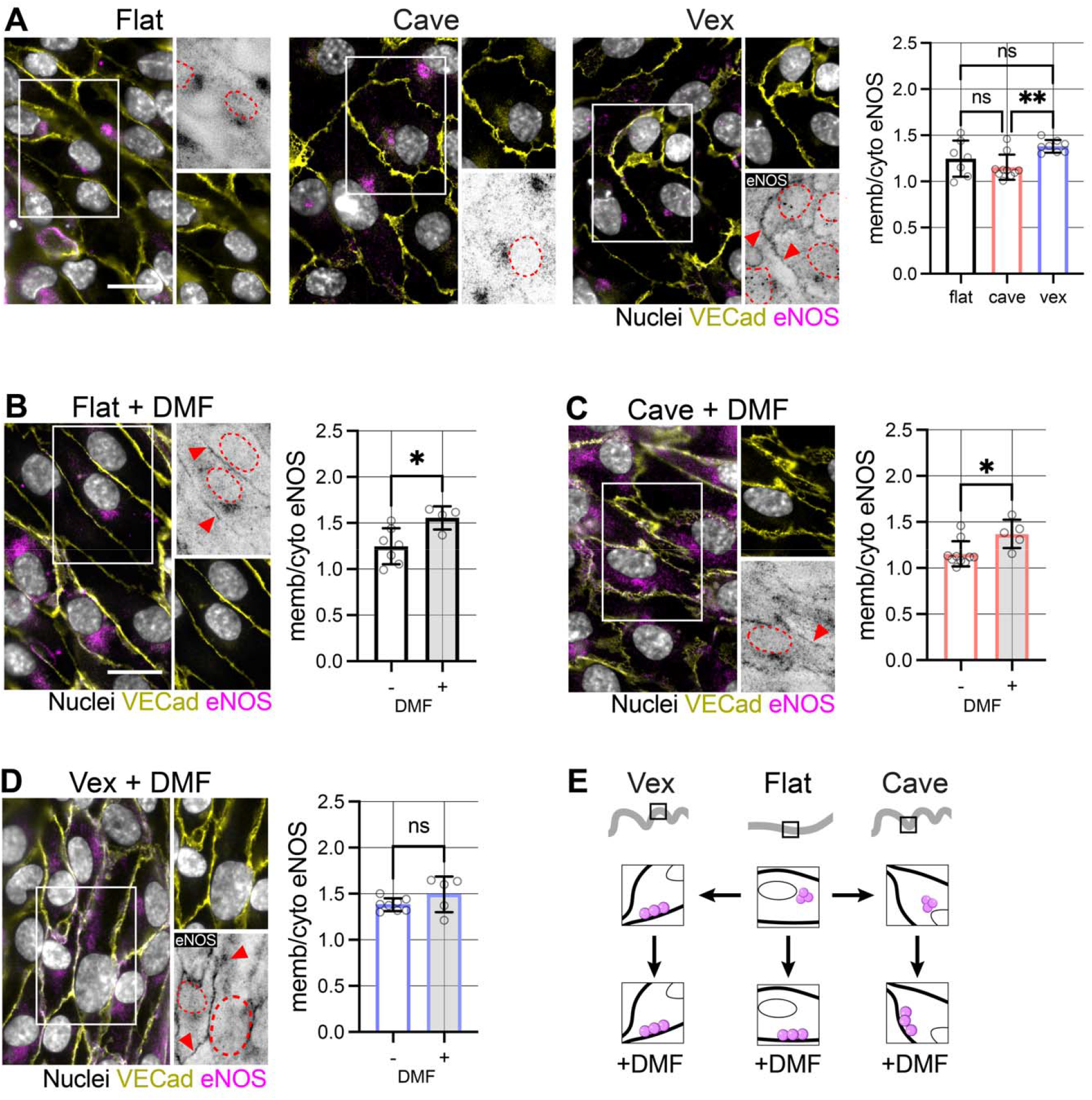
Convex curvature promotes plasma membrane localization of eNOS. **A** Representative fluorescent images with inverted grayscale insets of eNOS channel and quantification of membrane-to-cytoplasmic eNOS intensity ratio (memb/cyto eNOS) of hUVECs cultured atop programmed flat and TI 1.8 magnetic hydrogels with concave (cave) and convex (vex) regions (scale bars 20 μm, flat: n = 7 ROIs; each ROI = 100 μm^2^; N = 2, cave: n = 10 ROIs; each ROI = 100 μm^2^; N = 2, vex: n = 8 ROIs; each ROI = 100 μm^2^; N = 3). **B** Representative fluorescent images with inverted grayscale insets of eNOS channel and quantification of membrane-to-cytoplasmic eNOS intensity ratio (memb/cyto eNOS) eNOS of hUVECs cultured atop programmed flat NdFe-loaded DN hydrogels without ((−) DMF) and with dimethyl fumarate ((+)DMF) treatment at 24 h (scale bars 20 μm, (−)DMF: n = 7 ROIs; each ROI = 100 μm^2^; N = 2, (+)DMF: n = 4 ROIs; each ROI = 100 μm^2^; N = 2) **C** Representative fluorescent images with inverted grayscale insets of eNOS channel and quantification of membrane-to-cytoplasmic eNOS intensity ratio (memb/cyto eNOS) of hUVECs atop concave (cave) regions of TI 1.8 NdFe-loaded DN hydrogels without ((−) DMF) and with dimethyl fumarate ((+)DMF) treatment at 24 h (scale bars 20 μm, (−)DMF: n = 10 ROIs; each ROI = 100 μm^2^; N = 2, (+)DMF: n = 5 ROIs; each ROI = 100 μm^2^; N = 2). **D** Representative fluorescent images with inverted grayscale insets of eNOS channel and quantification of membrane-to-cytoplasmic eNOS intensity ratio (memb/cyto eNOS) of hUVECs atop convex (vex) regions of TI 1.8 NdFe-loaded DN hydrogels without ((−) DMF) and with dimethyl fumarate ((+)DMF) treatment at 24 h (scale bars 20 μm, (−)DMF: n = 8 ROIs; each ROI = 100 μm^2^; N = 3, (+)DMF: n = 5 ROIs; each ROI = 100 μm^2^; N = 2). **E** Schematic illustrating how sub-cellular localization of eNOS changes from being near the nucleus to near the cell membrane with development of convex curvature without (top) and with DMF treatment (bottom). **A-D** N = independent experiments; **p < 0.01, *p < 0.05, ns: not significant by two-tailed Student’s *t*-tests (two experimental groups) and one-way analysis of variance (ANOVA) with Bonferroni post hoc testing (more than two experimental groups).

To determine whether eNOS localization is regulated by YAP translocation, we treated cells with DMF at curvature induction and quantified eNOS distribution after 24 h. DMF increased the membrane-to-cytoplasmic eNOS ratio in both flat controls (Fig. 5B) and concave regions of TI 1.8 hydrogels (Fig. 5C) to the level of convex regions prior to DMF treatment (Fig. S5, Supporting Information). In contrast, DMF induced no further increase in the membrane-to-cytoplasmic eNOS ratio in convex regions (Fig. 5D), likely because it was already enriched at the plasma membrane. Together, these data indicate that convex curvature promotes membrane localization of eNOS, and that inhibition of YAP nuclear translocation induces a similar redistribution under conditions that exhibit predominantly membrane-associated eNOS (Fig. 5E).

## 3. Conclusion

In this study, we developed magnetically programmed hydrogels to dynamically introduce aspects of tortuous curvature into endothelial monolayers and isolate curvature as a mechanical cue independent of flow. While magnetic actuation of flat hydrogels increased their modulus, this change alone induced only limited endothelial responses. In contrast, dynamic curvature induced endothelial mechanical activation, marked by YAP nuclear localization that was strongest in regions of high convex curvature.

Curvature-induced remodeling was also reflected in endothelial function. Convex curvature promoted sustained thinning of VE-Cadherin junctions, consistent with junctional remodeling. In parallel, convex regions showed redistribution of eNOS towards the plasma membrane, indicating that local geometry influences endothelial functional state in addition to cytoskeletal and junctional remodeling. These findings suggest that vascular curvature may directly regulate endothelial function through local mechanical and topographical cues, even in the absence of secondary changes in flow. Importantly, many previous studies have been limited by curvature-associated changes in flow, which makes it difficult to interpret the independent contributions of geometry to endothelial cell function.[8]

More broadly, this work establishes a dynamic and tunable platform to study how evolving vascular geometry contributes to endothelial dysfunction. An important limitation of the current study is the lack of direct readouts of local barrier function, which remain challenging to obtain in a dynamic system. Future studies may address this through approaches such as immunofluorescence-based indirect permeability assays that allow fluorescence wherever there are discontinuities within the monolayer[32] to quantify local changes in barrier permeability. Similarly, although we observed curvature-dependent changes in intracellular eNOS localization, the present system does not yet allow direct correlation of these local changes with the nitric oxide output. Here, isolated convex and concave hydrogels[33] may provide a useful strategy to better define how eNOS localization relates to nitric oxide release[34,35].

Finally, although the present study used a 2.5D system with patterned concave and convex regions, future studies may extend this approach to 3D tubular geometries[36] and multicellular vascular models incorporating smooth muscle cells[37] or fibroblasts[38]. Such systems may provide a more complete framework for understanding how dynamic vessel remodeling contributes to vascular disease.

## 4. Experimental Section/ Methods

### Hydrogel preparation

#### Hydrogel Fabrication

Pre-hydrogel solution was prepared as previously described.[16] Briefly, 12 wt% acrylamide and 2 wt% alginate were dissolved in phosphate buffered saline (PBS). Dry Neodymium iron (NdFe) particles (Chemazone 5 μm, 918106-59-9) were then added at a 1:2 (wt./wt.) particle: pre-hydrogel ratio to form a slurry. Under constant stirring, crosslinkers and initiators were added to achieve final working concentrations of 0.012 wt.% N,N′-Methylenebisacrylamide, 2 mM ammonium persulfate, 0.165% tetramethylethylenediamine and 20 mM calcium sulfate. The pre-hydrogel solution was then pressed between Sigmacote^®^-treated coverslips, incubated at 50°C for 60 minutes, removed from the coverslips, and soaked overnight in PBS prior experimental use.

#### Programming Mold fabrication

The 3D mold designs were generated in Solidworks 2025 to program magnetic hydrogels with TI values of 1.0, 1.3, and 1.8. Molds were designed either as flat surfaces (TI 1.0) or with sinusoidal patterns of 0.32 mm (TI 1.3) and 0.69 mm (TI 1.8), each with a constant wavelength of 1 mm. Flat and patterned molds were then fabricated by 3D-printing using a Phrozen Sonic Mini 8K resin printer with Aqua Grey resin at a layer resolution of 50 μm.

#### Magnetic programming

Following overnight equilibrium swelling, magnetic hydrogels were cut into 10 × 8 mm rectangles and programmed by exposure to a 1 T magnetic field for 10 s while sandwiched between flat or patterned 3D-printed molds, yielding TI values of 1.0, 1.3, or 1.8. The required programming field was generated using the magneto-rheology accessory of an HR-30 Discovery Hybrid Rheometer (TA Instruments).

#### Magnetic actuation

Magnetized hydrogels were actuated by exposure to magnetic fields up to 60 mT generated by an electromagnet (Bunting BDE-4032-12) powered by a DC power supply (Tekpower TP3005P). For fields greater than 60 mT, cylindrical neodymium permanent magnets (McMaster-Carr, #5862K129) were positioned at varying distances from the hydrogels. Hydrogel dimensions during actuation were recorded using a Dino-Lite Premier AM3113 digital camera, and images were analyzed in FIJI (version: ImageJ 2.16).

#### Shear rheology

Magnetic hydrogels were crosslinked *in situ* between the plates of a temperature-controlled magneto-rheology accessory mounted on an HR-30 Discovery Hybrid Rheometer (TA Instruments) using a 20 mm, 2° cone angle geometry with a 57 μm gap. Shear storage and loss moduli were measured at room temperature at 1 Hz and 1% strain. The magneto-rheology accessory was used to apply magnetic fields as needed for shear measurements.

### Cell culture

#### Collagen coating of magnetic hydrogels

To functionalize hydrogels with collagen, 7-8 μL of 2 mg mL^−1^ *N*-Sulfosuccinimidyl-6-(4’-azido-2’-nitrophenylamino) hexanoate (Sulfo-SANPAH) in PBS was added to the surface of each magnetic hydrogel. Samples were then exposed to high-intensity UV light (37 mW cm^−2^, λ = 302 – 365 nm) for 5 min, followed by repeated PBS washes. Next, 0.25 mg mL^−1^ of bovine collagen type I (FibriCol^®^, Advanced Biomatrix, #5133) was added to the activated magnetic hydrogel surface and incubated overnight at 4 °C. The following day, excess collagen was removed by repeated PBS washes, and the hydrogels were sterilized under germicidal UV light for 1 h immediately prior to cell seeding.

#### hUVEC seeding and culture

hUVECs (Lonza - C2519A) were expanded in endothelial cell growth media (EBM-2 Basal Medium (CC-3156) supplemented with EGM^®^-2 SingleQuots^®^ (CC-4176), Lonza) and used up to passage 5. For seeding, cells were detached with 0.05% trypsin/EDTA for 8 min at 37°C to obtain a single cell suspension. Cells were counted, resuspended in fresh media, and seeded onto collagen-coated magnetic hydrogels at a density of 0.5 × 10^6^ per cm^2^. Cells were cultured for 5 days, with magnetic field exposure applied during the final 24 h of culture.

#### Small molecule inhibition

Perturbation studies targeting actin cytoskeletal organization and nuclear translocation of YAP were performed using 10 μM Y27632 (Tocris Bioscience, #1254251) and dimethyl fumarate (DMF, Sigma-Aldrich, #242926), separately added to the media on day 4 of hUVEC culture atop magnetic hydrogels.

### Staining and Imaging analyses

#### Cellular staining and immunofluorescence

Samples were fixed in 4% paraformaldehyde for 30 min at room temperature and washed three times with PBS. For immunofluorescence staining, samples were blocked in 2 wt% bovine serum albumin (BSA) for 1 hour at room temperature. YAP was labeled using a monoclonal anti-YAP antibody (1:200, Santa Cruz Biotech, #sc101199), and eNOS was labeled using a rabbit monoclonal antibody (Cell Signaling Technology, #32027) diluted in 2% BSA overnight at 4°C. Samples were then incubated for 1 h at room temperature in 2 wt% BSA containing the appropriate secondary antibody and counterstains: goat anti-rabbit IgG (1:200, Invitrogen, #A32731TR), goat-anti mouse IgG (1:200, Invitrogen, #A21235) phalloidin (1:250, Thermo Fisher Scientific, #A12380), and Hoechst (1:1000, Thermo Fisher Scientific, #62249). For VE-Cadherin staining, a directly conjugated VE-Cadherin Antibody (F-8) Alexa Fluor^®^ 488 (1:200, Santa Cruz Biotechnology, #sc-9989 AF488) was applied for 1 h at room temperature in 2 wt% BSA.

#### Imaging and analysis

All images were acquired using a Leica DMi8 THUNDER wide-field microscope with a 25x water-immersion objective, with 20-25% intensity and an exposure time of 200-260 ms. All imaging settings were kept consistent across experiments. For determining convex and concave regions on patterned samples, a low resolution (5x) image was taken, followed by at least 2 randomly chosen ROIs per concave and convex region. All image-based analyses were performed with ImageJ (U. S. National Institutes of Health, Bethesda, Maryland, USA, https://imagej.nih.gov/ij/). For YAP analysis, Otsu’s thresholding was applied to the Hoechst channel to define nuclei and to the YAP channel to define cellular area. The cytoplasmic mask was then generated by image subtraction. Mean YAP fluorescent intensity was measured in the nuclear and cytoplasmic compartments, and the nuclear-to-cytoplasmic intensity ratio was calculated for each cell which was averaged across each ROI. For VE-Cadherin junction thickness analysis, the VE-Cadherin channel was thresholded using Otsu’s method, and the binary mask was analyzed using the local thickness ImageJ plugin. This produced a false-color thickness map and a histogram of junction thickness values, from which the mean thickness was determined for each ROI. For eNOS membrane-to-cytoplasmic intensity analysis, a similar workflow as YAP was used, except that the membrane-to-cytoplasmic ratio was calculated for each cell and then averaged across each ROI.

#### Statistical analysis and reproducibility

Statistical analyses were performed using GraphPad Prism 9. Comparisons between two groups were conducted using two-tailed Student’s *t-*tests, whereas comparisons among multiple groups were performed using one-way ANOVA with Bonferroni post hoc correction. No data were excluded from the analyses. All experiments were performed in at least three independent replicates, as described in the text.

## Supporting Information

Supporting Information is available from the Wiley Online Library or from the author.

## Acknowledgements

This work was supported by funding from the NIGMS (R35 GM157063 to C.L.), the David and Lucile Packard Foundation (to C.L.), and the American Heart Association (24PRE1191531 to A.R.).

## References

[1] Loganathan, R. et al. (2020). Extracellular matrix dynamics in tubulogenesis. Cellular Signalling. 10.1016/j.cellsig.2020.109619.

[2] Smedby, Ö. and Bergstrand, L. (1996). Tortuosity and atherosclerosis in the femoral artery: What is cause and what is effect? Annals of Biomedical Engineering. 10.1007/BF02648109.

[3] Khosravani-Rudpishi, M. et al. (2018). The significant coronary tortuosity and atherosclerotic coronary artery disease; What is the relation? Journal of Cardiovascular and Thoracic Research. 10.15171/jcvtr.2018.36.

[4] Ekhator, C. et al. (2023). Arterial Tortuosity Syndrome: Unraveling a Rare Vascular Disorder. Cureus. 10.7759/cureus.44906.

[5] Kliś, K.M. et al. (2019). Tortuosity of the Internal Carotid Artery and Its Clinical Significance in the Development of Aneurysms. Journal of Clinical Medicine. 10.3390/jcm8020237.

[6] Zhang, X. et al. (2025). The relationship between retinal vascular tortuosity and retinal vasculitis. Journal of Ophthalmic Inflammation and Infection. 10.1186/s12348-025-00512-7.

[7] Katoh, K. (2023). Effects of Mechanical Stress on Endothelial Cells In Situ and In Vitro. International Journal of Molecular Sciences. 10.3390/ijms242216518.

[8] Mandrycky, C.J. et al. (2023). Endothelial Responses to Curvature-Induced Flow Patterns in Engineered Cerebral Aneurysms. Journal of Biomechanical Engineering. 10.1115/1.4054981.

[9] Chong, D.C. et al. (2017). Tortuous Microvessels Contribute to Wound Healing via Sprouting Angiogenesis. Arteriosclerosis, Thrombosis, and Vascular Biology. 10.1161/ATVBAHA.117.309993.

[10] Dessalles, C.A. et al. (2021). Integration of substrate- and flow-derived stresses in endothelial cell mechanobiology. Communications Biology. 10.1038/s42003-021-02285-w.

[11] Esch, M.B. et al. (2011). Characterization of In Vitro Endothelial Linings Grown Within Microfluidic Channels. Tissue Engineering Part A. 10.1089/ten.tea.2010.0371.

[12] Mannino, R.G. et al. (2015). “Do-it-yourself in vitro vasculature that recapitulates in vivo geometries for investigating endothelial-blood cell interactions.” Scientific Reports. 10.1038/srep12401.

[13] Paek, J. et al. (2021). Soft robotic constrictor for in vitro modeling of dynamic tissue compression. Scientific Reports. 10.1038/s41598-021-94769-2.

[14] Salipante, P.F. et al. (2022). Blood vessel-on-a-chip examines the biomechanics of microvasculature. Soft Matter. 10.1039/D1SM01312B.

[15] Davies, P.F. (2009). Hemodynamic shear stress and the endothelium in cardiovascular pathophysiology. Nature Clinical Practice Cardiovascular Medicine. 10.1038/ncpcardio1397.

[16] Roy, A. et al. (2024). Programmable Tissue Folding Patterns in Structured Hydrogels. Advanced Materials. 10.1002/adma.202300017.

[17] Park, J.W. et al. (2024). Development of stimuli-responsive flexible micropillar composites via magneto-induced injection molding and characterization of magnetic particle alignment. Polymer Testing. 10.1016/j.polymertesting.2023.108316.

[18] Swärd, K. et al. (2021). New Kids on the Block: The Emerging Role of YAP/TAZ in Vascular Cell Mechanotransduction, in Vascular Mechanobiology in Physiology and Disease, vol. 8, Springer International Publishing, Cham, pp. 69–96.

[19] Ritsvall, O. and Albinsson, S. (2024). Emerging role of YAP / TAZ in vascular mechanotransduction and disease. Microcirculation. 10.1111/micc.12838.

[20] Han, J. et al. (2025). Matrix Stiffness Regulates Mechanotransduction and Vascular Network Formation of hiPSC-Derived Endothelial Progenitors Encapsulated in 3D Hydrogels. 10.1101/2025.04.11.648340.

[21] Benson, J.C. and Brinjikji, W. (2020). The Chalice Sign: Characteristic Morphology of the Cervical Carotid Bifurcation in Patients with Loeys-Dietz Syndrome. Clinical Neuroradiology. 10.1007/s00062-019-00838-5.

[22] Doyle, M.G. et al. (2019). Comparison of Qualitative and Quantitative Assessments of Iliac Artery Tortuosity and Calcification. Vascular and Endovascular Surgery. 10.1177/1538574419858163.

[23] Ahmed, D.W. et al. (2025). Local photocrosslinking of native tissue matrix regulates lung epithelial cell mechanosensing and function. Nature Materials. 10.1038/s41563-025-02329-0.

[24] Giampietro, C. (2016). VE-cadherin complex plasticity: EPS8 and YAP play relay at adherens junctions. Tissue Barriers. 10.1080/21688370.2016.1232024.

[25] Khan, O.F. and Sefton, M.V. (2010). Perfusion and characterization of an endothelial cell-seeded modular tissue engineered construct formed in a microfluidic remodeling chamber. Biomaterials. 10.1016/j.biomaterials.2010.07.041.

[26] Birukov, K.G. et al. (2003). Magnitude-dependent regulation of pulmonary endothelial cell barrier function by cyclic stretch. American Journal of Physiology-Lung Cellular and Molecular Physiology. 10.1152/ajplung.00336.2002.

[27] Chien, S. (2008). Effects of Disturbed Flow on Endothelial Cells. Annals of Biomedical Engineering. 10.1007/s10439-007-9426-3.

[28] Van Epps, J.S. et al. (2009). Effects of Cyclic Flexure on Endothelial Permeability and Apoptosis in Arterial Segments Perfused Ex Vivo. Journal of Biomechanical Engineering. 10.1115/1.3192143.

[29] Fulton, D. et al. (2002). Localization of Endothelial Nitric-oxide Synthase Phosphorylated on Serine 1179 and Nitric Oxide in Golgi and Plasma Membrane Defines the Existence of Two Pools of Active Enzyme. Journal of Biological Chemistry. 10.1074/jbc.M106302200.

[30] Hu, Z. et al. (2013). Acute Mechanical Stretch Promotes eNOS Activation in Venous Endothelial Cells Mainly via PKA and Akt Pathways. PLoS ONE. 10.1371/journal.pone.0071359.

[31] Chen, F. et al. (2013). The Subcellular Compartmentalization of Arginine Metabolizing Enzymes and Their Role in Endothelial Dysfunction. Frontiers in Immunology. 10.3389/fimmu.2013.00184.

[32] Pramotton, F.M. et al. (2023). DYRK1B inhibition exerts senolytic effects on endothelial cells and rescues endothelial dysfunctions. Mechanisms of Ageing and Development. 10.1016/j.mad.2023.111836.

[33] Tomba, C. et al. (2022). Epithelial cells adapt to curvature induction via transient active osmotic swelling. Developmental Cell. 10.1016/j.devcel.2022.04.017.

[34] Villadangos, L. and Serrador, J.M. (2024). Subcellular Localization Guides eNOS Function. International Journal of Molecular Sciences. 10.3390/ijms252413402.

[35] Schilling, K. et al. (2006). Translocation of Endothelial Nitric-Oxide Synthase Involves a Ternary Complex with Caveolin-1 and NOSTRIN. Molecular Biology of the Cell. 10.1091/mbc.e05-08-0709.

[36] Son, J. et al. (2026). Triple□Scale Endothelialized Tubular Networks via Hybrid Biofabrication for Scalable Vascular Tissue Engineering. Advanced Healthcare Materials. 10.1002/adhm.202503334.

[37] Méndez-Barbero, N. et al. (2021). Cellular Crosstalk between Endothelial and Smooth Muscle Cells in Vascular Wall Remodeling. International Journal of Molecular Sciences. 10.3390/ijms22147284.

[38] Landau, S. et al. (2018). Oscillatory Strain Promotes Vessel Stabilization and Alignment through Fibroblast YAP□Mediated Mechanosensitivity. Advanced Science. 10.1002/advs.201800506.

