## Supplementary Information for "Magnetoactive hydrogels to probe curvature-directed endothelial cell mechanosensing"

A. Roy, C. Loebel,

Department of Materials Science & Engineering, University of Michigan, 2300 Hayward St., Ann Arbor, MI 48109-2117, USA

Center for Precision Engineering for Health (CPE4H), University of Pennsylvania, 25 N. 38th Street, Suite 700, Philadelphia, PA 19104, USA

R. Yanala,

Department of Materials Science and Engineering, University of Pennsylvania, 200 LRSM, 3231 Walnut St, Philadelphia, PA 19104-6272, USA

G. K. Hinds, J. YC. Liu, A. Velieva, C. Loebel

Department of Bioengineering, University of Pennsylvania, 240 Skirkanich Hall, 210 S 33rd St, Philadelphia, PA 19104-6321, USA

Center for Precision Engineering for Health (CPE4H), University of Pennsylvania, 25 N. 38th Street, Suite 700, Philadelphia, PA 19104, USA

**Contents**:

Figure S1: Shear storage modulus of programmed magnetic hydrogels

Figure S2: Regional YAP nucleus-to-cytoplasmic ratio after Y27 treatment

Figure S3. VE-Cadherin junction thickness after 24 h of curvature induction

Figure S4. VE-Cadherin junction thickness in DMF-treated cells

Figure S5. Membrane-to-cytoplasmic eNOS intensity ratio in DMF-treated cells

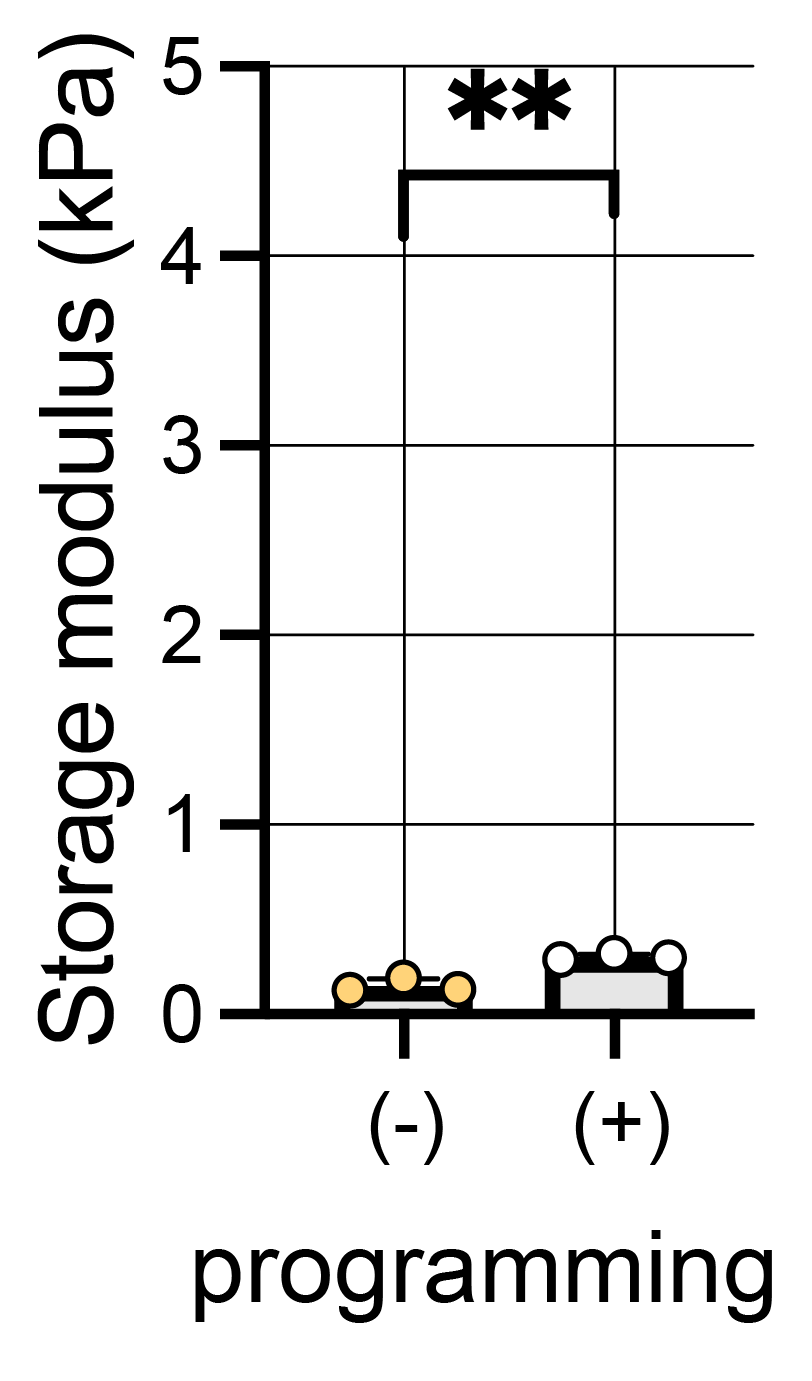

**Figure S1. Shear storage modulus of programmed magnetic hydrogels**

Quantification of shear storage modulus of magnetic hydrogels before and after programming with a 1 T magnetic field, measured in the absence of an actuation field. n = 3 hydrogels per group from 3 independent experiments; N = independent experiments, **p < 0.01, *p < 0.05, by two-tailed Student’s *t*-tests.

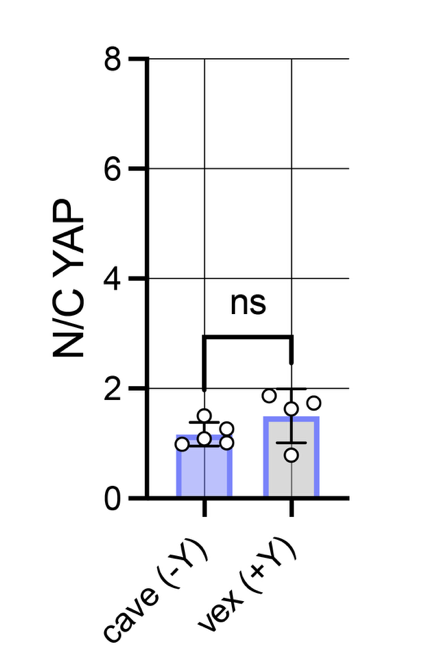

**Figure S2. Regional YAP nucleus-to-cytoplasmic ratio after Y27 treatment**

Quantification of nuclear-to-cytoplasmic (N/C) YAP intensity ratio in cells on concave (cave) regions of untreated (-Y) TI1.8 samples and cells on convex (vex) regions of TI1.8 samples treated with 10 μM Y27632 (+Y) at the onset of curvature induction. cave (-Y): n = 5 ROIs, N = 2; vex (+Y): n = 5 ROIs, N = 2; N = independent experiments, ns: not significant by two-tailed Student’s *t*-test

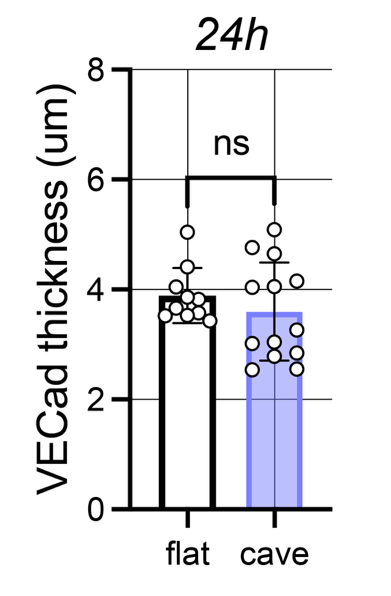

**Figure S3. VE-Cadherin junction thickness after 24 h of curvature induction.**

Quantification of VE-cadherin junction thickness in cells on flat samples and cells on concave (cave) regions of TI 1.8 samples after 24 h of curvature induction. flat: n = 10 ROIs, N = 3; cave: n = 13 ROIs, N = 3; N = independent experiments, ns: not significant by two-tailed Student’s *t*-test

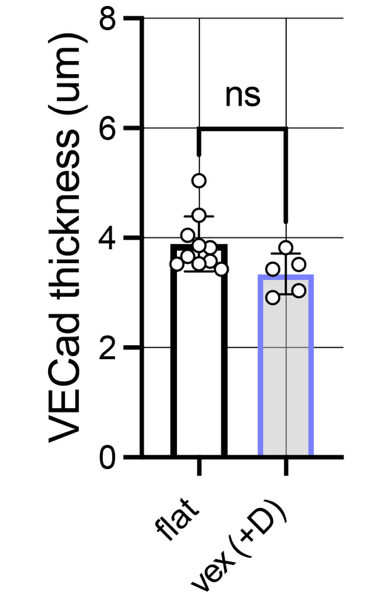

**Figure S4. VE-Cadherin junction thickness in DMF-treated cells**

Quantification of VE-cadherin junction thickness in cells on flat samples and cells on convex (vex) regions of TI1.8 samples treated with 50 μM DMF at the onset of curvature induction. flat: n = 10 ROIs, N = 3; vex (+D): n = 5 ROIs, N = 2; N = independent experiments, ns: not significant by two-tailed Student’s *t*-test

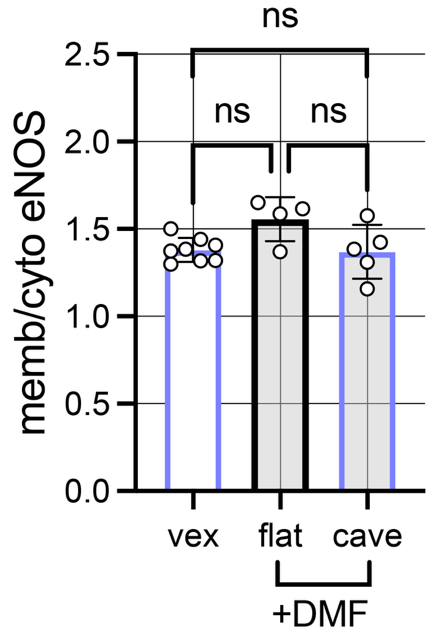

**Figure S5. Membrane-to-cytoplasmic eNOS intensity ratio in DMF-treated cells**

Quantification of membrane-to-cytoplasmic eNOS intensity ratio (memb/cyto eNOS) for cells on convex (vex) regions of untreated TI1.8 samples and for DMF-treated cells on flat and concave (cave) regions. vex: n = 8 ROIs, N = 3; flat (+D): n = 4 ROIs, N = 2; cave (+D): n = 5 ROIs, N = 2; N = independent experiments, ns: not significant by one-way ANOVA with Bonferroni post hoc correction.
